# Temporal Dynamics of BOLD fMRI Predict Intracranially-Confirmed Seizure Onset Zones in Drug-Resistant Epilepsy

**DOI:** 10.64898/2026.04.15.718821

**Authors:** Karl-Heinz Nenning, Erkam Zengin, Ting Xu, Elisabeth Freund, Noah Markowitz, Sarah Johnson, Silvia B Bonelli, Alexandre R. Franco, Stanley J. Colcombe, Michael P. Milham, Ashesh D. Mehta, Stephan Bickel

## Abstract

**Objective:** In individuals with drug-resistant epilepsy, accurately identifying the brain regions where seizures originate is a critical prerequisite to guide surgical treatment and achieve seizure freedom. To accomplish this, intracranial EEG is considered the gold standard, providing the spatiotemporal high-resolution data necessary to pinpoint epileptogenic activity. However, this precision is achieved through an invasive procedure with significant patient burden, which is fundamentally limited by the electrode placement and spatial coverage.

**Methods:** In this study, we investigated the potential utility of preoperative resting-state fMRI to non-invasively map alterations in brain dynamics at the whole brain level. Region-wise brain dynamics were quantified with complementary measures of local autocorrelation decay rates. We assessed the capacity of these derived features to effectively identify intracranial EEG confirmed seizure onset zones in 18 individuals with drug-resistant medial temporal lobe epilepsy. Overall, the study cohort contained 3867 implanted electrodes of which 159 classified as seizure onset zones by two independent board-certified epileptologists.

**Results:** Overall, our findings reveal more constrained temporal dynamics for brain regions associated with seizure onsets compared to non-seizure onset zones. Individual-level prediction showed a performance better than chance in 15 of the 18 patients. The overall predictive performance across all patients yielded a median AUC of 0.81, a median true positive rate of 0.75, and a median true negative rate of 0.83. Furthermore, in a subset of 13 patients, those with negative seizure outcomes showed higher probabilities of seizure onset zone predictions outside the resection area compared to those with good outcomes.

**Significance:** Overall, our findings suggest that altered temporal dynamics derived from preoperative resting-state fMRI represent a promising non-invasive approach for delineating epileptogenic tissue, potentially informing intervention strategies and guiding electrode placement.

## Introduction

For individuals with drug-resistant epilepsy, surgical resection or ablation is considered the best treatment option to achieve seizure freedom (de Tisi et al., 2011; Engel et al., 2012; Jobst & Cascino, 2015; Wiebe et al., 2001). A critical prerequisite for a successful treatment outcome is the accurate identification and delineation of the key hub of the epileptic network where seizures consistently originate, often termed the seizure onset zone (SOZ). To meet this clinical need, intracranial EEG (iEEG) is considered the gold standard for the definitive localization of the SOZ (Jayakar et al., 2016; Jobst et al., 2020; Rosenow & Lüders, 2001). By recording electrophysiological signals directly from the brain, iEEG provides the spatiotemporal high-resolution data necessary to pinpoint epileptogenic activity and guide surgical treatment. However, this precision is achieved through an invasive procedure with inherent risks and significant patient burden. Critically, the diagnostic power of iEEG is fundamentally limited by its restricted spatial coverage, introducing significant sampling bias and rendering interpretations highly dependent on electrode placement. This constraint makes the success of invasive iEEG examinations dependent on accurately targeting the electrode placement beforehand, a major challenge that, if unsuccessful, can lead to inconclusive results.

To improve surgery for individuals with drug-resistant epilepsy, it is therefore crucial to develop non-invasive tools that provide whole brain maps of functional features that facilitate pinpointing exactly where seizures originate (Corona et al., 2023; Sathe et al., 2023; Stufflebeam et al., 2011). Here, leveraging brain dynamics is particularly valuable as subtle, pathological alterations in the temporal structure of brain activity may be present even during the interictal state (Fisher et al., 2014; Gerstl et al., 2023; Lai et al., 2023), potentially reflecting underlying alterations specific to epileptogenic regions. Also, epilepsy has been increasingly recognized as a network disease with dysfunctions beyond a focal seizure onset, with regions showing interictal abnormalities being part of the wider epileptogenic network (Bernhardt et al., 2015; Burns et al., 2014; Kanner et al., 2017; Kramer & Cash, 2012; Larivière et al., 2022; Piper et al., 2022; Schaper et al., 2023; Scharfman et al., 2018). Analyzing the temporal dynamics of brain activity by utilizing non-invasive techniques with a high spatial coverage, such as functional magnetic resonance imaging (fMRI), could potentially be used to identify unique functional characteristics of the seizure onset zone and its brain-wide interactions (Richardson, 2012). Critically, non-invasive whole brain mapping of functional abnormalities could provide a more comprehensive view of the epileptogenic network, thereby improving the selection of candidates for surgery and guiding the placement of intracranial electrodes.

In this study, we investigate the potential utility of preoperative fMRI to non-invasively map alterations in brain dynamics in epilepsy patients with medial temporal lobe seizure onset. We assess the capacity of these observed brain dynamics to effectively identify iEEG-confirmed seizure onset zones on the individual patient-level.

## Materials and Methods

### Study Cohort

We studied 18 individuals with drug-resistant epilepsy and medial temporal lobe seizure onset (10 left, 6 right, 2 bilateral). All individuals underwent evaluation for epilepsy surgery at Northwell Health, NY, including magnetic resonance imaging (MRI) at 3-Tesla with qualitative assessment by expert neuroradiologists for structural abnormalities. Individuals underwent stereoelectroencephalography (SEEG) to identify epileptic seizure origins. The seizure onset zones were localized from prolonged interictal and ictal monitoring data by two independent board-certified epileptologists. Overall, the study cohort contained 3867 implanted electrodes of which 159 were identified as seizure onset channels. All patients provided informed written consent according to a protocol approved by the institutional review board at the Feinstein Institutes for Medical Research in accordance with the Declaration of Helsinki. To establish a homogeneous study cohort, patients were included with SEEG confirmed TLE, and no other significant brain abnormality. Patient demographics are given in Table 1. As a control group, we utilized data from a group of 652 healthy controls (mean age 54.8 years, 330 female) based on the CamCAN study (Shafto et al., 2014; Taylor et al., 2017) and obtained from the CamCAN repository (https://cam-can.mrc-cbu.cam.ac.uk/dataset/).

| ID | Sex | Age | SOZ | ILAE (follow-up) | Resection | Electrodes (SOZ confirmed) |
| --- | --- | --- | --- | --- | --- | --- |
| P01 | F | 16 | left | 4 (3 years) | yes | 74 (7) |
| P02 | F | 46 | right | 1 (2 years) | yes | 240 (7) |
| P03 | F | 36 | left | 1 (2 years) | yes | 205 (4) |
| P04 | M | 43 | left | 4 (3 years) | yes | 162 (5) |
| P05 | M | 47 | right | n/a | n/a | 234 (11) |
| P06 | F | 30 | bilateral | 3 (2 years) | yes | 245 (15) |
| P07 | F | 37 | left | 5 (1 years) | yes | 175 (5) |
| P08 | M | 22 | right | 4 (2 years) | yes | 217 (4) |
| P09 | M | 46 | left | 4 (1 years) | n/a | 306 (9) |
| P10 | F | 32 | left | 4 (1 years) | yes | 336 (7) |
| P11 | F | 22 | right | 3 (1 years) | yes | 150 (13) |
| P12 | F | 29 | left | 1 (2 years) | yes | 300 (5) |
| P13 | F | 27 | left | 5 (2 years) | yes | 187 (6) |
| P14 | F | 24 | left | 1 (1 years) | yes | 148 (11) |
| P15 | F | 59 | right | n/a | n/a | 181 (2) |
| P16 | M | 21 | left | 1 (1 years) | yes | 115 (9) |
| P17 | M | 20 | right | 1 (3 years) | n/a | 240 (8) |
| P18 | M | 21 | bilateral | 4 (3 years) | n/a | 352 (31) |

### MRI data acquisitions

MRI data was acquired using a 3-Tesla Siemens Prisma MRI, equipped with a 32-channel head coil. The imaging session comprised acquisition of a high resolution structural T1-weighted MPRAGE volume (TR/TE 2400/2.2 ms, flip angle 8°, 208 × 300 × 320 matrix, voxel size 0.8 × 0.8 × 0.8 mm). 5 patients (P01-P05) had one functional MRI sequence (TR/TE 2000/30 ms, flip angle 7°, 64 × 64 × 32 matrix, voxel size 3.75 × 3.75 × 4 mm, 240 volumes, 8 minutes), and 13 patients had two repeated runs of with a multiband sequence (TR/TE 720/33 ms, flip angle 52°, 104 × 90 × 72 matrix, voxel size 2.2 × 2.2 × 2 mm, multiband acceleration factor: 8, 594 volumes, 7.1 minutes) with anterior-posterior (run-1) and posterior-anterior (run-2) phase encodings.

### Electrode localization

Electrode locations were identified and visualized using the iELVis toolbox (Groppe et al., 2017). Briefly, the preoperative T1-weighted anatomical scan was used to perform tissue segmentation and pial surface reconstruction with FreeSurfer (Dale et al., 1999; Fischl et al., 1999). Then, postoperative CT scans were acquired and coregistered to the FreeSurfer reconstruction. Contacts were then semi-manually localized using the BioImage Suite software (version 3.01) (Papademetris et al., 2006). This established electrode locations (coordinates) in native space.

### Imaging data preprocessing

Minimal image preprocessing was performed with fMRIPrep (v22.1.1) (Esteban et al., 2018). In brief, the anatomical T1 volumes were skull-stripped, segmented into brain tissues of cerebrospinal fluid (CSF), white matter (WM) and gray matter (GM), and spatially normalized to the MNI standard space (template MNI152NLin6Asym). Resting-state fMRI data was first motion corrected, slice-timing corrected, and co-registered to the anatomical T1 volume. Subsequently, the fMRI volumes were normalized to MNI space by applying the warp obtained from the high-resolution anatomical scan and resampled to 2mm isotropic resolution. The fMRI data was denoised by regressing out confounding artifacts based on the head motion parameters and anatomical CompCor, and a bandpass filter (0.01-0.1Hz) was applied. Finally, the fMRI data was smoothed with at 4mm full-width at half-maximum, and voxel-wise time-series were extracted based on a gray matter segmentation of the MNI152NLin6Asym template. Where available, structural volumes of post-operative imaging data were used to manually segment the extent of the resection.

### Quantifying temporal dynamics

We measured local neural dynamics at the voxel-level with four established complementary methods to quantify intrinsic timescales, utilizing the temporal autocorrelation function (ACF) of resting-state brain activity. Together, they provide a fingerprint of temporal dynamics, where shorter timescales relate to faster changes in brain activity and longer timescales reflect more slower, constrained functional dynamics. Specifically, we quantified the following timescale variants. To obtain a more holistic characterization of the ACF decay, we quantified the sum of the values in the first positive period of the autocorrelation function (ints_00) (Watanabe et al., 2019). To capture distinctions between the steepness of the ACF decay, we established the coefficients (τ) of an exponential curve fit 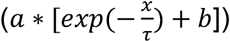 to the ACF (ints_exp) (Ito et al., 2020; Murray et al., 2014). Additionally, we also quantified the time for the ACF to decay to half (ints_05) (Raut et al., 2020), and the autocorrelation value at lag 1 (ints_t1). Features were extracted at the voxel level for each patient and the cohort of healthy controls. For each ACF decay measure and voxel, we quantified the mean and standard deviation in the healthy controls as a normative baseline. For each patient, the timescale measures were projected into the control baseline distribution by subtraction of its mean and division by its standard deviation, resulting in a z-score that characterizes the deviation from the controls. A schematic of the workflow is displayed in Figure 1.

**Figure 1.**
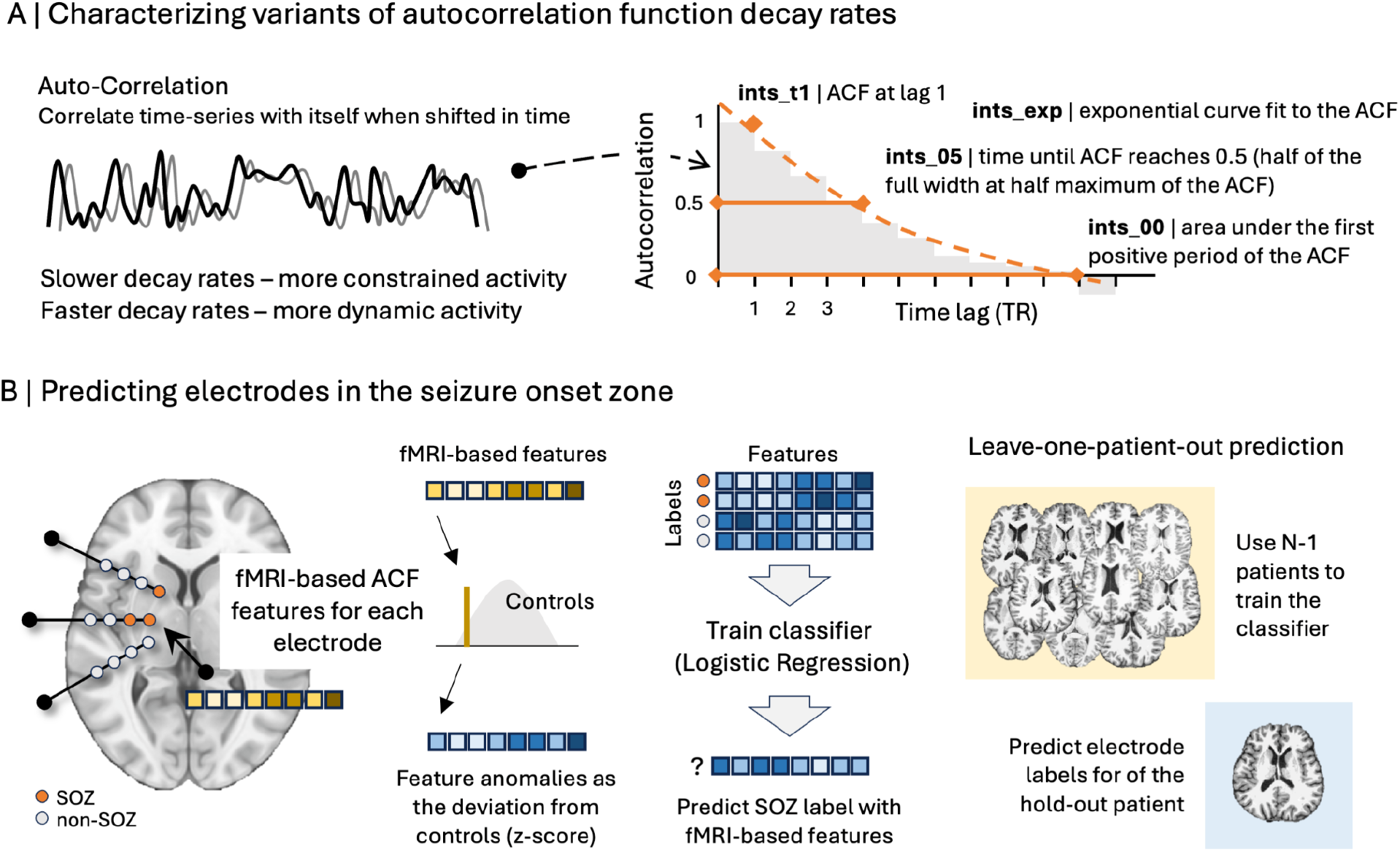
Schematic workflow. A) Variants of autocorrelation decay rate measures characterize different dynamic properties of the time-series dynamics. B) The seizure onset zone (SOZ) prediction workflow.

### Predictive Modeling

For each electrode, the clinical classification into (non-)seizure onset zones was used as binary class labels. A feature vector was established for each electrode by extracting the average value of each calculated feature in a sphere with a 3mm diameter around the electrode location. The features and labels were used in a binary classification scheme, using logistic regression with a l2 regularization and balanced classes. In a leave-one-patient-out cross-validation scheme, we trained the classifier on N-1 patients and predicted the electrode labels for each fMRI run of the held-out patient individually. Predictive performance was assessed with the area under the receiver operator curves (AUC). To quantify true positive and false negative rates, patient specific classification thresholds were established with the optimal Youden’s index (Youden, 1950). To detect potential overfitting and to exclude that predictions were generated by chance, 1,000 control runs with randomly shuffled electrode labels were performed.

## Results

### Seizure onset zone has altered autocorrelation decay rates during resting-state fMRI

Brain regions located within the seizure onset zones showed resting-state fMRI dynamics that differed from non-SOZ regions in three out of four autocorrelation decay measures (Figure 2A). A more constrained activity (slower temporal autocorrelation decay rates) was found for the coefficients of an exponential curve fit to the ACF (ints_exp; Mann-Whitney U test, p_FDR_=0.0004, rank-biserial correlation effect size 0.13), the area under the first positive period of the autocorrelation function (ints_00; Mann-Whitney U test, p_FDR_=2.13e-06, rank-biserial correlation effect size=0.17), and the autocorrelation value at lag 1 (ints_t1; Mann-Whitney U test, p_FDR_=6.01e-15, rank-biserial correlation effect size=0.28).

**Figure 2.**
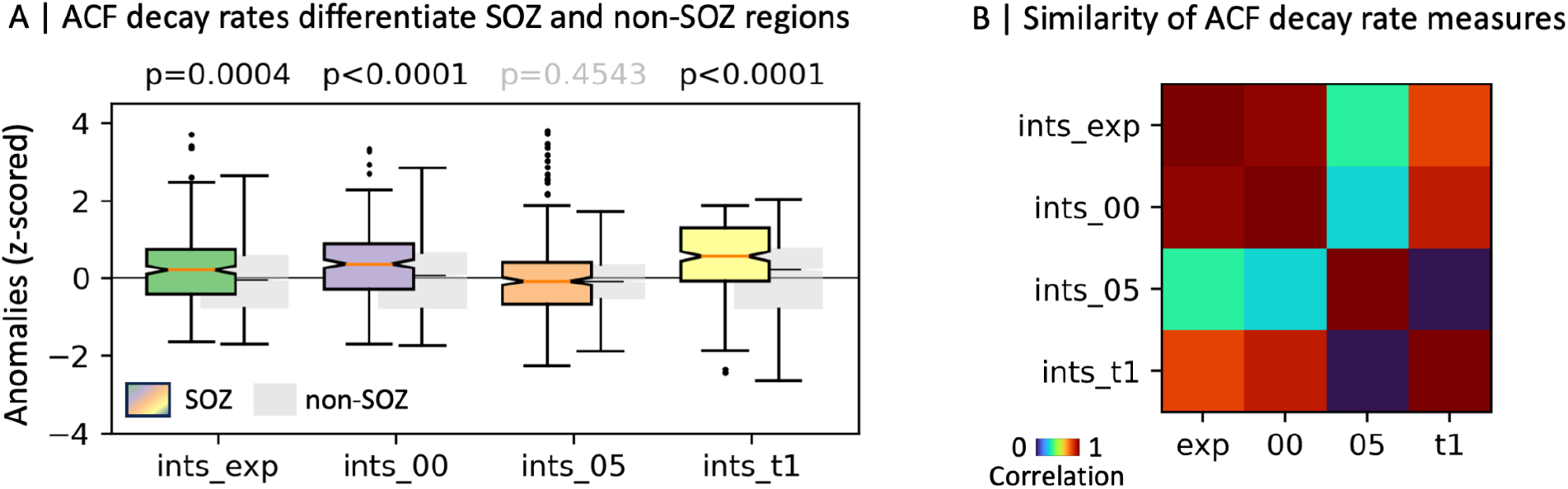
Comparison between the different autocorrelation function (ACF) decay rate features. A) Compared to electrodes located in non-epileptogenic tissue, the electrodes in seizure onset zones (SOZ) show divergent resting-state fMRI dynamics in three out of four ACF features. Longer timescales (decreased autocorrelation decay rates) were observed for coefficients of an exponential curve fit to the ACF (ints_exp), the area under the first positive period of the autocorrelation function (ints_00), and the autocorrelation value at lag 1 (ints_t1). The time for the ACF to decay to half (ints_05) did not show any differences between regions located inside and outside the seizure onset zones (Mann-Whitney U test, fdr corrected). B) ACF measures that quantify prolonged decay characteristics (ints_exp and ints_00) were the most similar. The autocorrelation half-life (ints_05), the time it takes for the signal’s correlation to drop by half, was the most distinct from the other measures.

Among the ACF measures that quantify prolonged decay characteristics (Figure 2B), ints_exp and ints_00 were the most similar to each other (r=0.97). The autocorrelation half-life (ints_05), the time it takes for the signal’s correlation to drop by half, was the most distinct from the other measures (mean r=0.23). The autocorrelation value at lag 1 (ints_t1) was highly similar to the coefficients of an exponential curve fit to the ACF (ints_exp) and the area under the first positive period of the autocorrelation function (ints_00) (mean r=0.87), but not to the autocorrelation half-life (ints_05) (r=0.02).

### fMRI-based autocorrelation features can predict electrodes located in the seizure onset zone

In a leave-one-patient-out scheme, the ACF decay rate measures showed promising results for predicting the electrodes that were iEEG-confirmed seizure origins. In 15 out of the 18 patients, we observed a predictive performance better than chance (Figure 3A). Within these patients, both repeated resting-state scans showed a comparable predictive performance. In three patients we observed poor predictions (Figure 3B). Overall, AUC rates were promising (median=0.81), and better than chance (paired t-test vs chance: t=8.86, p=7.15e-10, Cohen’s d=2.29). Binary classifications with patient-specific classification thresholds yielded a median true positive rate of 0.75 (paired t-test vs chance: t=9.05, p=4.38e-10, Cohen’s d=2.28) and a median true negative rate of 0.83 (paired t-test vs chance: t=5.71, p=3.12e-06, Cohen’s d=1.42) (Figure 3C). The coefficients of the logistic regression showed that the features based on the autocorrelation value at lag 1 (ints_t1) consistently have the highest weights of the logistic regression model (Figure 3D).

**Figure 3.**
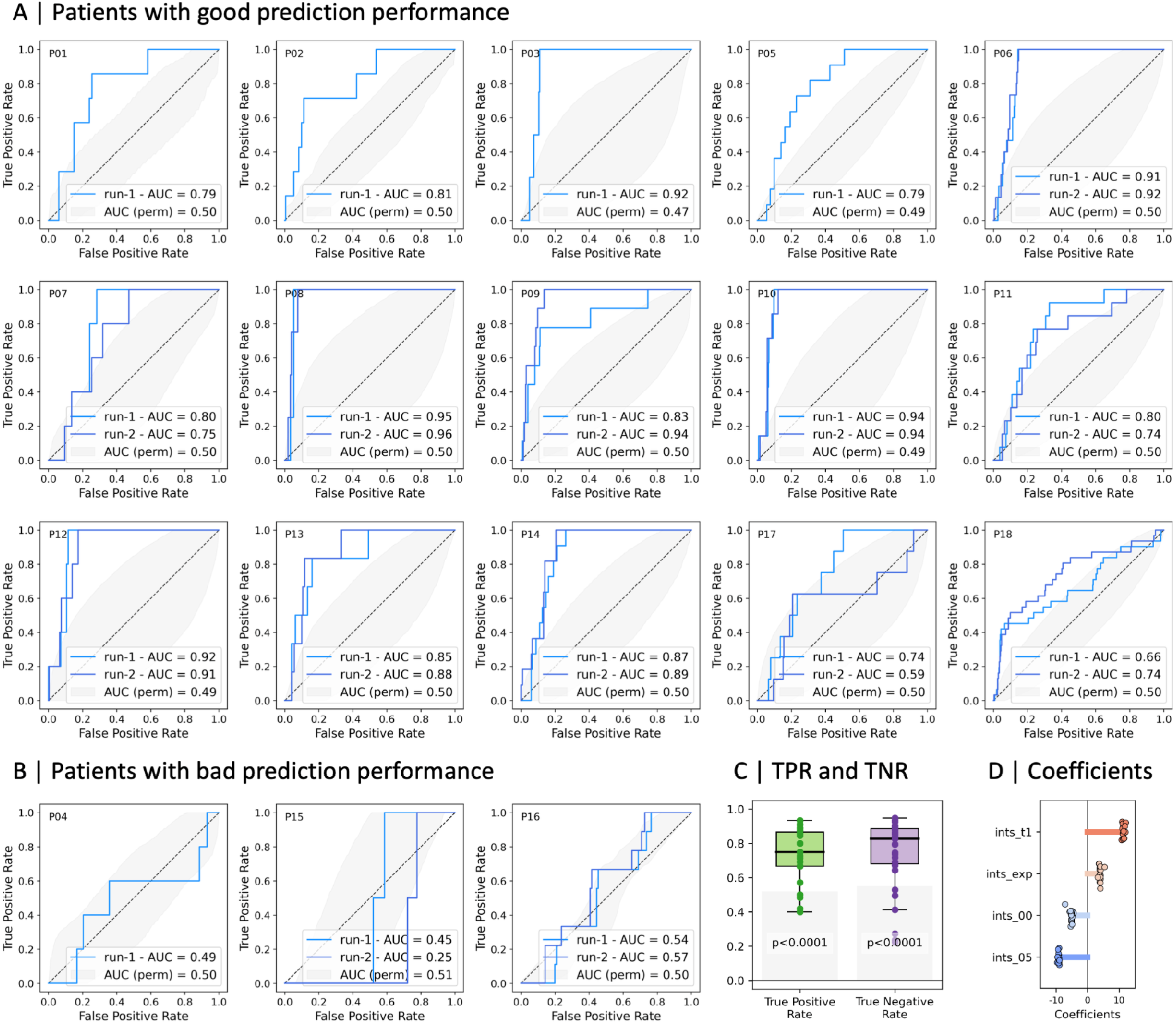
Prediction results of the leave-one-patient-out scheme. A) In 15 out of 18 patients a prediction performance better than chance was achieved. B) Three out of the 18 patients showed poor prediction performance. C) Overall, the area under the curve (AUC), and true positive (TPR) and negative (TNR) rates were better than chance. D) The coefficients of the logistic regression show that the features based on the autocorrelation value at lag 1 (ints_t1) have consistently the highest weights of the prediction model.

Our cohort was restricted to patients with a medial temporal lobe seizure onset, which was reflected by the localization of electrodes where the seizure onset labels were correctly (true positives) and falsely (false positives) predicted (Figure 4). Electrodes with a false positive prediction were located mainly in the vicinity of the seizure onset zones in the medial temporal lobe. Electrodes with true negative prediction results were located more laterally relative to the medial cortex, farther from the true seizure onset sites.

**Figure 4.**
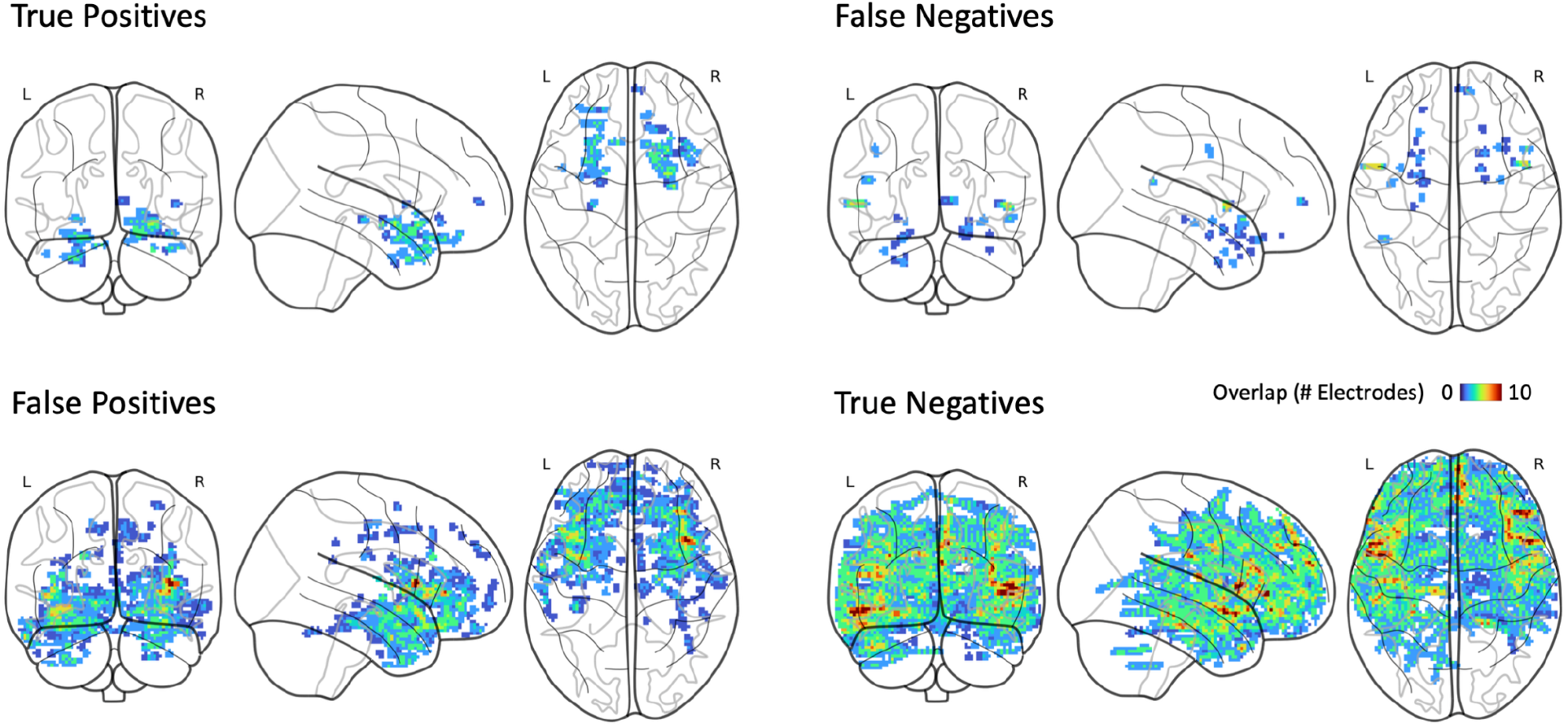
Electrodes with a false positive prediction were located in the vicinity of the seizure onset zones in the medial temporal lobe, and true negative predicted electrodes were located farther from the true seizure onset sites.

### False positive prediction results are associated with seizure outcome

In a subset of patients (n=13) with available imaging and outcome data post-surgery (ILAE 1: n=5; ILAE 3: n=2; ILAE 4: n=4; ILAE 5: n=2), we examined the relationship between prediction results and resection (Figure 5). Electrodes located within the resection zone had higher prediction probability in the patients with a good outcome (ILAE 1) than in patients with a worse outcome (ILAE 3, 4, and 5) (Kolmogorov-Smirnov test, KS statistic=0.53, p=0.0001). In contrast, electrodes outside the resection zone were more likely to be predicted as seizure onset zone in patients with a negative outcome (Kolmogorov-Smirnov test, KS statistic=0.16, p=4.44e-16).

**Figure 5.**
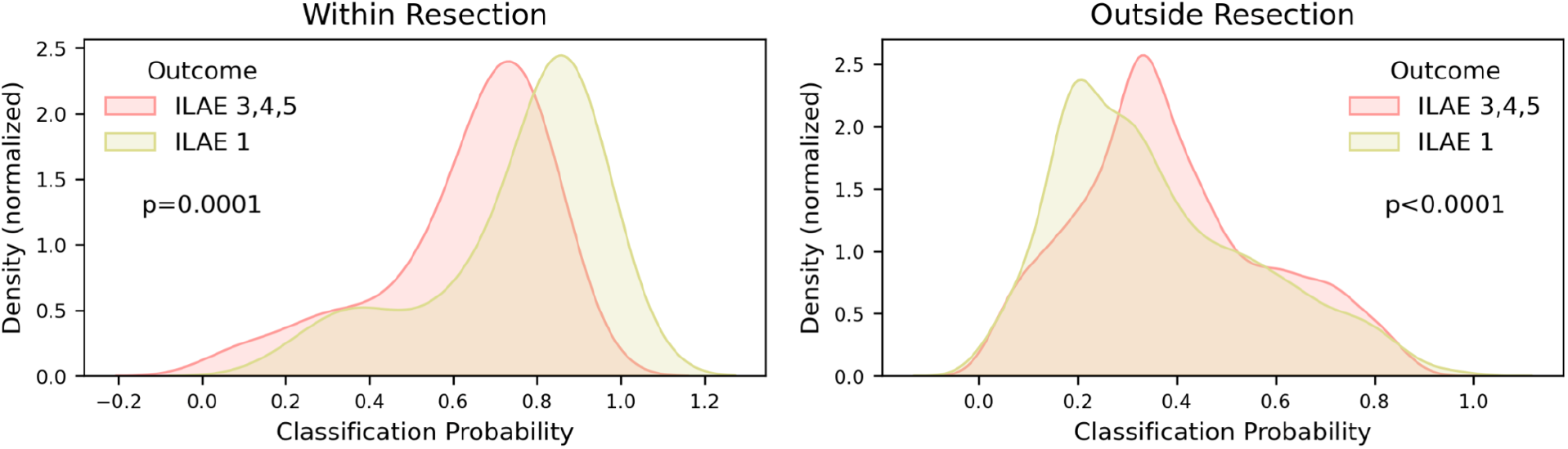
Association between prediction probabilities and outcome in a subset of patients (n=13) with available imaging and outcome data post surgery (ILAE1: n=5, ILAE 3: n=2, ILAE 4: n=4, and ILAE 5: n=2). In patients with a worse outcome (ILAE 3, 4, and 5), higher prediction probabilities were observed for electrodes outside of the resection zone.

## Discussion

This study investigated the utility of preoperative resting-state fMRI for predicting iEEG-identified seizure onset zones in patients with drug-resistant medial temporal lobe epilepsy. Region-wise brain dynamics were quantified with complementary measures of autocorrelation decay rates, characterizing temporal dependencies in brain activation. A leave-one-patient-out cross-validation framework was used to predict electrode labels (SOZ vs. non-SOZ) across the study population of 18 patients. At the individual level, prediction results better than chance were achieved in 15 out of the evaluated 18 patients. The overall predictive performance across all patients yielded a median AUC of 0.81, a median true positive rate of 0.75, and a median true negative rate of 0.83. These findings suggest that altered temporal dynamics derived from preoperative resting-state fMRI represent a promising non-invasive approach for delineating potential seizure onset zones.

### Local resting-state fMRI dynamics characterize the seizure onset zone

While epileptic seizures are defined as transient events with distinct boundaries (Fisher et al., 2005, 2017), abnormal interictal neuronal activity represents a hallmark of epilepsy, critically informing diagnosis and treatment strategies (Pillai & Sperling, 2006). Consequently, interictal EEG monitoring is an essential part of clinical care in epilepsy, providing information about the diverse nature of brain dynamics at a high temporal resolution (Fisher et al., 2014; Gerstl et al., 2023; Lai et al., 2023). Our findings extend this understanding by suggesting that altered interictal brain dynamics, in particular within brain regions associated with seizure onset, can also be observed with resting-state fMRI at much lower temporal scales. Indeed, resting-state fMRI has been repeatedly shown to capture meaningful brain characteristics in individuals with epilepsy (Bernhardt et al., 2016; Cataldi et al., 2013; Lucas et al., 2023; Pressl et al., 2018; Reyes et al., 2016), and it has been successfully employed to establish associations between altered functional connectivity patterns and potential seizure onset locations (Sathe et al., 2023; Stufflebeam et al., 2011). While early fMRI research heavily relied on static connectivity analyses, which provided valuable insights into average network interactions, more recent studies increasingly focus on brain dynamics to understand the temporal fluctuations of brain activity in epilepsy (Abreu et al., 2019; F. Liu et al., 2017; Pang et al., 2021).

One prominent measure of such temporal dynamics are so-called intrinsic timescales, reflecting temporal dependencies and the duration over which neural activity is sustained or integrated (Golesorkhi et al., 2021; Ito et al., 2020; Murray et al., 2014). By adopting this dynamic approach, our study utilized previously established and complementary measures of intrinsic timescales based on autocorrelation decay rates (Ito et al., 2020; Murray et al., 2014; Raut et al., 2020; Watanabe et al., 2019). Recent studies examining such timescales in epilepsy have shown varying findings. Specifically, widespread increased decay rates (shorter timescales) were observed in TLE when using the area under the ACF as a measure (Xie et al., 2023). In contrast, an exponential fit to the ACF revealed more constrained dynamics with slower decay rates (longer timescales) specifically proximal to the presumed epileptogenic zone (Nedic et al., 2015), and single-voxel autocorrelation over a fixed window of time also showed longer hippocampal timescales in left TLE (Bouffard et al., 2026). These findings are not necessarily contradicting, as different cortical regions can exhibit different autocorrelation properties that might not be fully captured with individual measures (Ito et al., 2020; Raut et al., 2020). Here, we used a complementary set of previously reported timescale measures to address this limitation and to account for nuances of the autocorrelation decay, such as the steepness of the initial decay, a flattening of decay rates, or zero crossings.

Overall, our findings indicate slower decay rates, characteristic for more restricted temporal dynamics, specifically in brain regions identified as seizure onset. This could indicate a tendency for neural activity in the epileptogenic tissue to persist longer in specific states, potentially reflecting underlying pathological hypersynchronization or a reduced capacity for efficient information processing compared to unaffected brain tissue (Clawson et al., 2023; Engel, 1996; Ferrario & Giovagnoli, 2023; Lemieux et al., 2011; S. Liu & Parvizi, 2019; Proix et al., 2017). The fact that our results corroborate previous localized findings in patients with epilepsy (Bouffard et al., 2026; Nedic et al., 2015) strengthens the evidence that altered intrinsic timescales are indeed a robust feature of the epileptogenic cortex.

Intriguingly, this interictal state of slower temporal dynamics resonates with the concept of ‘critical slowing down’ in dynamical systems. This phenomenon, characterized by a system taking longer to return to its baseline state after minor perturbations, is considered an indicator for the approaching of a critical transition point (Scheffer et al., 2009). Specifically in epilepsy, critical slowing down has been observed in intracranial EEG signals preceding seizure onset and interpreted as evidence of the brain transitioning towards the ictal state (Jirsa et al., 2014; Litt et al., 2001; Lopes da Silva et al., 2003; Maturana et al., 2020; McSharry et al., 2003; Moraes et al., 2021). Our resting-state fMRI findings during the interictal period raise the compelling possibility that SOZ regions inherently exist in a state closer to such a critical threshold. The underlying reduced interictal dynamic range might reflect tissue properties that render these areas more susceptible to the perturbations that can trigger a transition into a seizure. Therefore, quantifying these slower intrinsic timescales using non-invasive resting-state fMRI could offer a valuable pathophysiological marker, potentially aiding the clinical challenge of delineating epileptogenic tissue and identifying regions with heightened seizure risk to improve diagnostic accuracy and surgical outcomes.

### The spatial pattern of false positives may reflect broader network alterations

While the true positive rates for predicting electrodes in the seizure onset zone are encouraging, the concurrent observation of a substantial number of false positives warrants careful consideration. Critically, these false positive predictions were not randomly distributed across the brain, but instead primarily located within the medial cortex, anatomically close to the medial temporal lobe SOZs identified by intracranial EEG. The spatial clustering of these misclassifications near the actual SOZ, rather than in more remote lateral areas, suggests that our prediction model, based on resting-state fMRI temporal dynamics, is indeed sensitive to specific pathophysiological signatures related to epileptogenicity, even if these signatures might extend beyond the clinician-defined onset zones.

This pattern aligns with the current understanding of epilepsy, which is increasingly recognized as a dysfunction within broader brain networks - a network disease (Bernhardt et al., 2015; Burns et al., 2014; Kanner et al., 2017; Kramer & Cash, 2012; Larivière et al., 2022; Piper et al., 2022; Schaper et al., 2023; Scharfman et al., 2018). The network perspective emphasizes that even seizures originating focally rapidly engage and are sustained by interconnected brain regions, forming an epileptogenic network. This network encompasses not only the SOZ but also areas crucial for seizure propagation and regions exhibiting interictal abnormalities that might reflect underlying vulnerability or functional coupling to the primary focus. Therefore, it is plausible that the false positive regions identified by our model represent components of this wider epileptogenic network (Berg et al., 2010). Although these areas may not exhibit the earliest electrographic seizure onset activity captured by iEEG, they might still possess altered intrinsic temporal dynamics similar to those detected in the SOZ due to shared pathological influences, functional connectivity, or involvement in interictal discharge propagation (Boddeti et al., 2025; Ji et al., 2025; Larivière et al., 2021; Wang et al., 2023). Consequently, while technically false positives relative to the stringent iEEG definition of the SOZ, these spatially related predictions could hold valuable clinical information, potentially mapping the broader functional territory of the epileptogenic network involved in an individual patient’s epilepsy. Intriguingly, we observed higher probabilities for SOZ predictions outside of the resection zone in patients with a negative seizure outcome (ILAE 3, 4, and 5) than in patients with a good outcome (ILAE 1). This could indicate an unsuccessful resection of a wider epileptogenic network responsible for the bad outcome, potentially due to suboptimal electrode placement.

### Clinical utility of resting-state fMRI dynamics in presurgical planning

Despite its inherent risks and patient burden, invasive intracranial EEG monitoring is often a crucial step for the accurate localization of the seizure onset zone in drug-resistant epilepsy. Our findings demonstrate that altered temporal dynamics quantified with non-invasive, preoperative resting-state fMRI are associated with the iEEG-confirmed SOZ, highlighting a potential clinical utility of this technique. Specifically, leveraging pre-operative neuroimaging data for fMRI-guided placement of iEEG electrodes poses a promising application, potentially optimizing implantation strategies through indicating regions with aberrant functional dynamics. By better targeting electrodes towards areas exhibiting these functional signatures of epileptogenicity, resting-state fMRI could potentially increase the diagnostic yield of iEEG examinations or perhaps even allow for effective monitoring with reduced electrode coverage in certain patients, thereby minimizing procedural risks.

Furthermore, preoperative measures of resting-state fMRI dynamics could serve as an important component in the multi-modal process of generating hypotheses about SOZ location before invasive monitoring. When integrated with evidence from scalp EEG, structural MRI lesion analysis, or PET metabolism, the distinct patterns of fMRI dynamics associated with the SOZ could add a unique layer of converging evidence, helping clinicians refine their localization hypotheses (Corona et al., 2023; Lundstrom et al., 2021; Sathe et al., 2023; Stufflebeam et al., 2011). While the ultimate goal of replacing iEEG with purely non-invasive SOZ prediction requires significant further development, validation in larger and more diverse patient cohorts, and likely integration of multiple biomarkers, our results contribute to the growing evidence that resting-state fMRI dynamics offer valuable, clinically relevant information (Jiang et al., 2022). Given the broad application of pre-operative neuroimaging assessment, this approach holds considerable promise for complementing existing diagnostic tools and potentially streamlining the presurgical pathway by improving the accuracy and efficiency of SOZ localization.

### Limitations

Several limitations of this study should be considered when interpreting the results. First, a fundamental challenge for comparing invasive and non-invasive modalities is the discrepancy in spatial resolution. While iEEG records from a precise, millimeter-scale source, the fMRI BOLD signal is coarser, and our use of 4mm smoothing likely introduces partial volume effects, causing electrode signals to include contributions from adjacent tissue. This spatial mixing complicates the interpretation of false positives near the seizure onset zone, making it difficult to disentangle signal leakage from genuine pathological alterations in the bordering network. However, the fact that true negatives were predominantly located in lateral regions distal to the seizure onset suggests that our approach reliably identifies healthy dynamics when spatially distinct. This capacity to rule out unaffected regions could guide electrode placement and help optimize the targeting of invasive monitoring.

Second, imaging acquisition parameters and scan duration can influence the stability of resting-state fMRI measures, and the optimal sequence parameters for autocorrelation decay measures in epilepsy remain an open question. Notably, although our study cohort included scans with differing repetition times, the normalization of patient features against a normative control baseline effectively bridged these acquisition differences, suggesting robustness across sequence parameters. Although we observed consistent predictive performance across repeated runs in the majority of patients, longer acquisition times could potentially reduce noise and improve the reliability of the calculated timescales.

Finally, our sample size (N=18) and the restriction to patients with medial temporal lobe epilepsy limit the generalizability of our findings. While focusing on a homogeneous pathology allowed us to reduce biological variance, it remains to be determined whether these specific alterations in autocorrelation decay rates extend to extratemporal or neocortical focal epilepsies. Furthermore, while the use of a large, external healthy control dataset provided a robust normative baseline, differences in scanner hardware and demographics between the clinical and control sites cannot be fully ruled out as confounding factors. Validation in larger, multi-center cohorts with age-matched, site-specific controls will be essential to establish the clinical utility of this approach.

## Conclusion

In conclusion, our findings reveal more constrained temporal dynamics for brain regions associated with the seizure onset zone compared to non-SOZ areas. These results demonstrate the promise of using resting-state fMRI-derived temporal dynamics as a non-invasive tool to help delineate epileptogenic tissue, potentially informing intervention strategies. While false positive classification rates highlight the need for further evaluation, the observed widespread alterations emphasize the notion of epilepsy as a network disorder affecting brain regions beyond the focal onset site. Future research correlating these dynamic fMRI markers with detailed electrophysiological data and long-term patient outcomes is essential to clarify whether these false positives represent elements of the broader epileptogenic network (such as propagation pathways or irritable cortex), functional deficit zones, or other phenomena, thereby refining the specificity and clinical applicability of this approach for presurgical planning.

## Acknowledgements

Data collection and sharing of the healthy control group used in this project was provided by the Cambridge Centre for Ageing and Neuroscience (CamCAN). CamCAN funding was provided by the UK Biotechnology and Biological Sciences Research Council (grant number BB/H008217/1), together with support from the UK Medical Research Council and University of Cambridge, UK.

## Data availability statement

No data are available due to patient confidentiality issues.

## Ethics approval statement

This study was approved by the institutional review board at the Feinstein Institutes for Medical Research in accordance with the Declaration of Helsinki.

## Patient consent statement

All patients provided informed written consent.

